# Edlib: a C/C++ library for fast, exact sequence alignment using edit distance

**DOI:** 10.1101/070649

**Authors:** Martin Šošić, Mile Šikić

## Abstract

We present Edlib, an open-source C/C++ library for exact pairwise sequence alignment using edit distance. We compare Edlib to other libraries and show that it is the fastest while not lacking in functionality, and can also easily handle very large sequences. Being easy to use, flexible, fast and low on memory usage, we expect it to be a cornerstone for many future bioinformatics tools.

Source code, installation instructions and test data are freely available for download at https://github.com/Martinsos/edlib, implemented in C/C++ and supported on Linux, MS Windows, and Mac OS.

## 1 Introduction

One of the fundamental operations in bioinformatics is pairwise sequence alignment - a way to measure either the similarity or distance between two sequences. Due to the quadratic time complexity, deterministic algorithms that yield optimal alignment are inefficient for the comparison of long sequences. Therefore, they are used in the very last step when the aligning substrings of the given sequences are roughly determined using heuristic methods.

Deterministic, optimal alignment algorithms are unavoidable for the resequencing of genomes when the exact alignments of reads and reference are necessary for the successful determination of differences – especially the existence of single nucleotide variants. Owing to that, many aligners use some of the efficient variants of these algorithms for the final phase. For example, SNAP (Zaharia et al., 2011) uses Landau-Vishkin (Landau et al., 1986) as the core component.

The increased need for exact algorithms that could align longer segments has recently emerged as a consequence of the advent of long-read sequencing technologies such as Pacific Biosciences Single Molecule Real-Time (SMRT) sequencing technology and Oxford Nanopore Technologies (ONT), which produce reads over 10 kbp in length.

Deterministic methods can be categorized as local, global, or semi-global (overlap) alignment methods, regarding their scoring scheme. The basic global alignment algorithm is the dynamic programming Needleman-Wunsch algorithm (Needleman and Wunsch, 1970), and the basic local alignment algorithm is its variation, the Smith-Waterman algorithm (Smith and Waterman, 1981). Semi-global alignment methods, quite popular for read alignment, are similar to global alignment but they do not penalize gaps at the beginning or/and the end of the sequences. Both Needleman-Wunsch and Smith-Waterman algorithms have quadratic time and space complexity, so there has been a lot of work on trying to improve that. Ukkonen's banded algorithm (Ukkonen, 1985) reduces needed time by cleverly reducing the space of search, while Hirschberg's algorithm (Hirschberg, 1975) trades space for speed, reducing space complexity from quadratic to linear.

An important sub-category of alignment methods is the calculation of Levenshtein distance - the minimum number of single-character edits (insertions, deletions or substitutions) required to change one sequence into the other (also referred as edit distance). Myers managed to exploit its special properties by developing a bit-vector algorithm (Myers, 1999), reducing the computation time by a constant factor. Although it is one of the fastest deterministic alignment algorithms it is quite complex to implement and does not support global alignment. Hence, it is rarely implemented in practice. In this article, we present Edlib - our implementation of Myers's bit-vector algorithm, extended with additional methods and features that are important for its practical application.

## 2 Methods

Myers's bit-vector algorithm transforms dynamic programming matrix so that each cell can have only a binary value, which enables us to store multiple cells as a bit-vector into one CPU register and achieve single instruction multiple data (SIMD) parallelization. Myers additionally applies Ukkonen's banded algorithm to reduce the number of calculated bit-vectors.

The original algorithm was designed only for a semi-global alignment method where gaps at the start and at the end of the query sequence are not penalized (we named this method HW). In Edlib, we extended Myers's algorithm to support the global alignment method and the semi-global alignment method where gaps at the end of the query sequence are not penalized (we named this method SHW). For this, we came up with extended banded algorithm that also supports SHW and global method (Supplementary Methods). Unlike the original Myers's implementation, it is not necessary to define the size of the band in advance. The algorithm begins with a narrow band and widens it until find an optimal alignment.

Originally, Myers's algorithm returns no information about the optimal alignment path – the optimal sequence of edit operations that need to be performed on the query sequence to transform it to the target sequence. In Edlib, we further extended Myers's algorithm with the finding of the optimal alignment path for all three supported alignment methods in linear space by combining it with Hirschberg's algorithm. Inspired by the SSW Library (Zhao, 2013), we reduced a problem of finding the path for HW and SHW to finding the path for global alignment, which both simplifies the implementation and improves the computation speed (Supplementary figure 1).

We implemented Edlib as both C/C++ library and a stand-alone application. Supported operating systems are MS Windows, Linux, and Mac OS.

## 3 Results

We compared Edlib to SeqAn library (Döring et al., 2008), Parasail library (Daily, 2016) and original Myers's implementation.

We chose SeqAn to be the center of our comparison since it is one of the most advanced sequence alignment libraries currently available and the only C/C++ library that has all the methods supported by Edlib. Döring et al. (2008) show that SeqAn is the library with the fastest implementation of sequence alignment using edit distance (they also combine Myers's bit vector algorithm with Hirschberg's algorithm), and to our knowledge, there are no developments up to now showing otherwise.

Comparison was done against SeqAn v2.1.1 and Parasail v1.0.3, which were the latest releases at the moment of writing this article.

The tests were performed on Linux, Intel Core i7-4710HQ 2.5 Ghz with 16GB RAM. As test data, we used real DNA sequences ranging from 10-5000 kbp in length and their artificially mutated versions, in order to show how the similarity and length of aligned sequences affect performance.

The run times for finding the global alignment edit distance (no alignment path) are displayed in Table 1. Results show that Edlib is 2.5-250 times faster than SeqAn, and 12-3500 times faster than Parasail, the difference being the largest when the sequences are large and similar. Additionally, we ran similar tests (Supplementary Table 1, Supplementary Table 2, Supplementary Table 3) for the semi-global methods where Edlib also outperformed both SeqAn and Myers's implementation, while Parasail does not support the semi-global methods. Finally, we also ran a test for the finding of alignment path (Supplementary table 2), however, SeqAn failed to produce results, and Parasail and Myers do not support the finding of alignment path.

**Table 1.** Edlib, SeqAn, and Parasail run time comparison of finding global alignment edit distance for different sequence lengths and similarities.

| Seq. sizes | Similarity | **Edlib** | SeqAn | Parasail |
| --- | --- | --- | --- | --- |
| $10^6 \times 10^6$ | 99.3% | **0.4s** | 112s | 1397s |
| $10^6 \times 10^6$ | 92.5% | **8.5s** | 112s | 1419s |
| $10^6 \times 10^6$ | 81.3% | **15s** | 112s | 1330s |
| $10^6 \times 10^6$ | 64.0% | **32s** | 112s | 1459s |
| $10^6 \times 10^6$ | 51.1% | **29s** | 112s | 1461s |
| $10^5 \times 10^5$ | 99.3% | **0.011s** | 1s | 4.8s |
| $10^5 \times 10^5$ | 92.9% | **0.044s** | 1s | 4.8s |
| $10^5 \times 10^5$ | 82.4% | **0.21s** | 1s | 4.8s |
| $10^5 \times 10^5$ | 66.2% | **0.43s** | 1s | 4.8s |
| $10^5 \times 10^5$ | 53.9% | **0.4s** | 1s | 4.8s |

As can be seen from the results, Edlib exhibits significant improvement in speed with increase of sequence similarity, in contrast to other libraries. This is due to our implementation of the banded algorithm, which significantly reduces search space for similar sequences.

The similarity of two sequences was calculated as 1 − *edit_distance / min(length_query_, length_target_*). Two different DNA sequences were used for these tests. We artificially mutated them to achieve different similarities. Myers's implementation is not included in this comparison as it does not support global alignment.

## Acknowledgements

The authors would like to thank Ivan Sović for valuable help with testing Edlib and providing comments on the manuscript.

